# Improving protein function prediction with synthetic feature samples created by generative adversarial networks

**DOI:** 10.1101/730143

**Authors:** Cen Wan, David T. Jones

## Abstract

Protein function prediction is a challenging but important task in bioinformatics. Many prediction methods have been developed, but are still limited by the bottleneck on training sample quantity. Therefore, it is valuable to develop a data augmentation method that can generate high-quality synthetic samples to further improve the accuracy of prediction methods. In this work, we propose a novel generative adversarial networks-based method, namely FFPred-GAN, to accurately learn the high-dimensional distributions of protein sequence-based biophysical features and also generate high-quality synthetic protein feature samples. The experimental results suggest that the synthetic protein feature samples are successful in improving the prediction accuracy for all three domains of the Gene Ontology through augmentation of the original training protein feature samples.

## Introduction

Protein function prediction is an important but challenging task in bioinformatics. The challenge comes from the inherent high dimensionality of the input feature space and the cryptic relationship between sequence and function. The importance is clear from the fact that very few proteins in data banks have complete or reliable functional annotations. Up to the year 2015, fewer than 0.1% of proteins deposited in the UniProt database had received even just one experiment-based functional annotation, and less than 20% of proteins had been even just electronically annotated in all three domains of gene function as defined by the Gene Ontology [1]. Although recent community-wide efforts have overall pushed forward the development of computational prediction methods, the accuracy of predicting the vast majority of protein functions is still very low [2] [3] [4]. This is not only due to the fact of the natural diversity of protein function, but also because of the limited number of existing functionally annotated protein samples. This issue leads to a bottleneck in the performance of prediction methods, especially for machine learning-based methods [5], [6], [7], on making accurate predictions based on such small reference or training datasets. Due to the expense of obtaining protein function data experimentally, it is highly desirable to develop computational methods that can make better use of existing limited data. To that end, here we explore the possibility that high-quality synthetic samples can be created to augment the existing annotation data and further improve the predictive accuracy of our prediction models.

Generative Adversarial Networks (GANs) [8] [9] [10] [11] [12] [13] are a new type of generative model, which aim to generate high-quality synthetic samples by accurately learning the underlying distributions of target data samples. The novel aspect of GANs is that they adopt an adversarial training paradigm, where two neural networks “fight” against each other to learn the distribution of samples. One network (the generator) attempts to generate synthetic data, and the other network (the discriminator) attempts to decide whether a given sample is real or synthetic. Each network gets better and better at its task until an equilibrium is reached, where the generator can’t make better samples, and the discriminator can’t detect more synthetic samples. GANs have already shown outstanding performance on different machine learning tasks in the image processing field, such as image to image translation [14] [15] [16], image segmentation [17] [18] [19] and image reconstruction [20] [21] [22]. In addition to handling image data, GANs have also performed well with other types of data, such as gene expression data and raw gene sequence data. Wang, et. al (2018) [23] and Dizaji, et. al (2018) [24] proposed a conditional GAN-based framework for the task of gene expression profiles inference by modeling the conditional distribution of target genes given the corresponding landmark genes’ profiles. Ghahramani, et al. (2018) [25] also proposed a WGAN-GP-based method to capture the diversity of cell types based on the large and sparse scRNA-seq data. More recently, Gupta & Zou (2019) [26] and Wang, et. al (2019) [27] successfully proposed GAN-based methods to generate synthetic genes and promoters, respectively.

The data augmentation task is also an area where GANs show a great potential to achieve good performance. Most of the existing work on GAN-based data augmentation methods also focus on the image processing tasks, such as image classification [28] [29] [30]. For example, Frid-Adar, et. al (2018) [28] adopted the well-known DCGAN [9] method to generate synthetic liver lesion images, which successfully improved the accuracy of liver lesion classification. Most recently, Marouf, et. al (2018) [31] adopted GANs to generate synthetic scRNA-seq profiles, which were used for downstream cell type classification tasks. In this work, we propose a new GAN-based data augmentation approach – FFPred-GAN, which is the first work up to the present that successfully employs GANs to cope with protein sequence-based data distributions to tackle the protein function prediction problem. More specifically, the novelties are three-fold. To begin with, FFPred-GAN successfully learns the distribution of protein amino acid sequence-based biophysical features and generates high-quality synthetic protein feature samples. Moreover, those high-quality synthetic protein feature samples successfully augment the original training samples and obtain significantly higher accuracy in predicting all three domains of GO terms. Furthermore, FFPred-GAN also shows good computational time efficiency, which is valuable when dealing with the large amount of sequence data in present data banks. These properties also encourage further extension of FFPred-GAN in exploiting other types of protein-related features.

## Results

### Overview of FFPred-GAN

In general, the FFPred-GAN framework consists of three steps to generate high-quality synthetic training protein feature samples, as shown in Figure 1. To begin with, FFPred-GAN adopts the widely-used FFPred [32] feature extractor to derive protein biophysical information based on the raw amino acid sequences. For each input protein sequence, 258 dimensional features are generated to describe 13 groups of protein biophysical information, such as secondary structure, amino acid composition and presence of motifs. Then FFPred-GAN adopts the Wasserstein Generative Adversarial Network with Gradient Penalty (WGAN-GP) approach [11] to learn the actual high-dimensional distributions of these training proteins’ features. The generator of WGAN-GP is used to output the synthetic training protein feature samples during different training stages of FFPred-GAN. On the last step, FFPred-GAN uses the Classifier Two-Sample Tests (CTST) [33] to select the optimal synthetic training protein feature samples, which are used to augment the original training samples. During the down-stream machine learning classifier training stage, the optimal synthetic samples are expected to derive better classifiers, leading to higher predictive accuracy.

**Figure 1:**
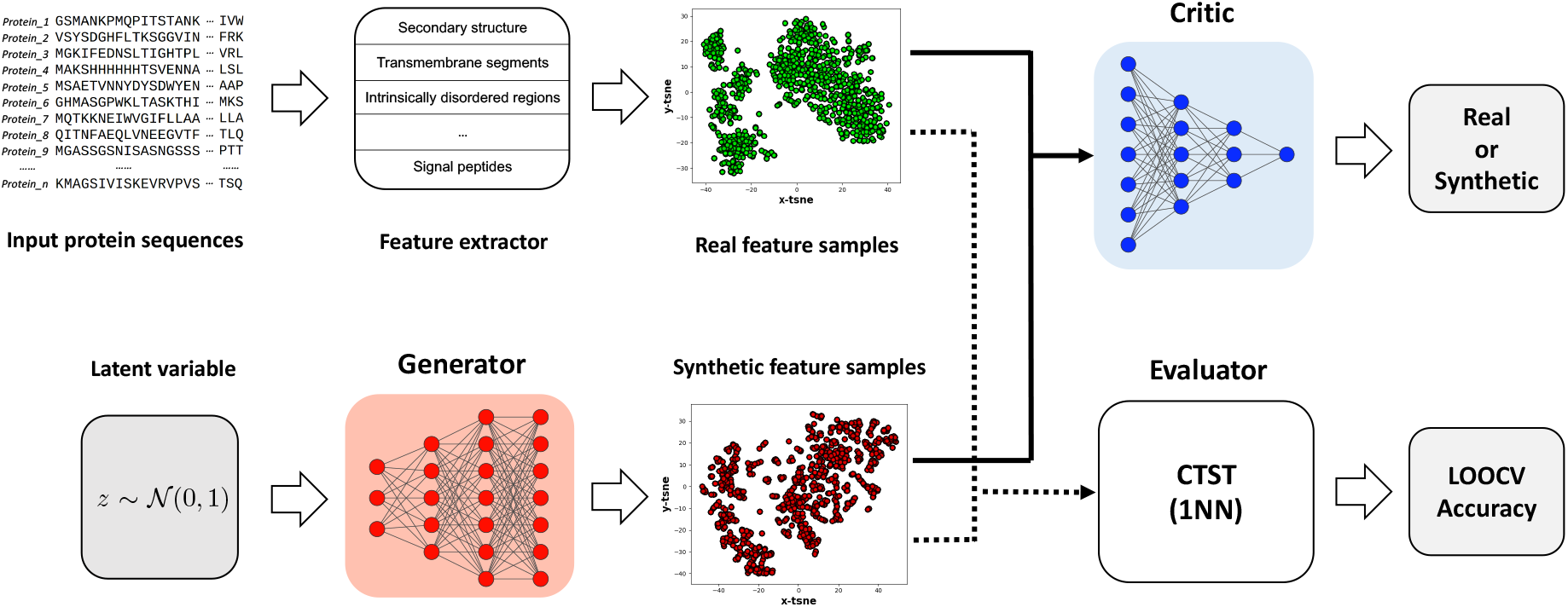
The flowchart of FFPred-GAN.

### FFPred-GAN successfully generates high-quality synthetic protein biophysical feature samples

In general, FFPred-GAN successfully learns the distributions of the training protein biophysical feature samples and generates high-quality synthetic ones. We train two FFPred-GAN models for each GO term by using two different sets of protein samples with different class labels. The first FFPred-GAN model is trained by using the protein samples that are annotated with that GO term (hereafter we denote those proteins as the so-called positive samples). The other FFPred-GAN model is trained by using the protein samples that are not annotated by that GO term (hereafter we denote those proteins as the so-called negative samples). Therefore, in total, we train 602 FFPred-GAN models for all 301 GO terms in the FFPred-fly library.

As mentioned in the last section, we adopt the 1-nearest neighbour classification algorithm and the Leave One Out Cross-Validation (LOOCV) to conduct the classifier-two-sample tests, which is used for evaluating the quality of synthetic protein feature samples. The closer the value of LOOCV accuracy is to 0.500, the higher the quality of synthetic samples. As shown in Figures 2.A – 2.C, the star and circle symbols respectively denote the LOOCV accuracies obtained for synthetic positive and negative protein feature samples generated by individual GO term-based FFPred-GANs. The *x*-axis denotes the indexes of GO terms, while the *y*-axis denotes the LOOCV accuracy that ranges from 0.000 to 1.000. In general, the synthetic positive protein feature samples generated by FFPred-GAN for nearly half of BP, MF and CC terms obtained a LOOCV accuracy of 0.500. The average LOOCV accuracies for the BP, MF and CC domains of GO terms are 0.573, 0.584 and 0.590, respectively. Figures 2.D – 2.H display the *t*-SNE transformed 2D visualisation of real and synthetic positive protein feature samples that are generated during different training stages of FFPred-GAN for the BP term GO:0000375. In detail, at the begin of FFPred-GAN training (i.e. after the 1^*st*^ epoch), the real positive protein feature samples (green dots) are distributed distantly from the synthetic ones (red dots), leading to a LOOCV accuracy of 1.000, suggesting obvious differences between the real and synthetic sets of protein feature samples. After 1,000 epochs of further training, FFPred-GAN shows that it has started capturing the distribution of the real protein feature samples and generating synthetic ones that are starting to be similar to the real ones, due to the fact that the distributions of the red and green dots have overlapping areas around the diagonal with a better LOOCV accuracy of 0.737. After even further training of FFPred-GAN, on the 10,001^*st*^ epoch, the overlapping areas of two sets of protein samples become broader, leading to the LOOCV accuracy of 0.645, which also indicates significantly improved training quality of FFPred-GAN. The training quality of FFPred-GAN continues to improve with more epochs of training, with the LOOCV accuracy reaching 0.515 after another 10,000 epochs. Finally, after 29,601 epochs’ training, FFPred-GAN has been successfully trained due to the desired LOOCV accuracy of 0.500. Also, as shown in Figure 2.H, both sets of protein feature samples project to almost the exact same areas. This pattern is consistent for training FFPred-GAN for the positive protein feature samples for the MF and CC domains of GO terms. As shown in Figures S1.A – S1.E and S1.F – S1.J respectively, the quality of GO:0000981 and GO:0000785’s synthetic positive protein feature samples gradually becomes better with the increasing number of training epochs. The green and red dots for the synthetic positive protein feature samples of GO:0000981 are distributed similarly after 44,201 epochs training. Analogously, the distributions of synthetic positive protein feature samples for GO:0000785 also becomes non-significantly different to the corresponding real one after 31,801 epochs training, since the LOOCV accuracy reaches to 0.500.

**Figure 2:**
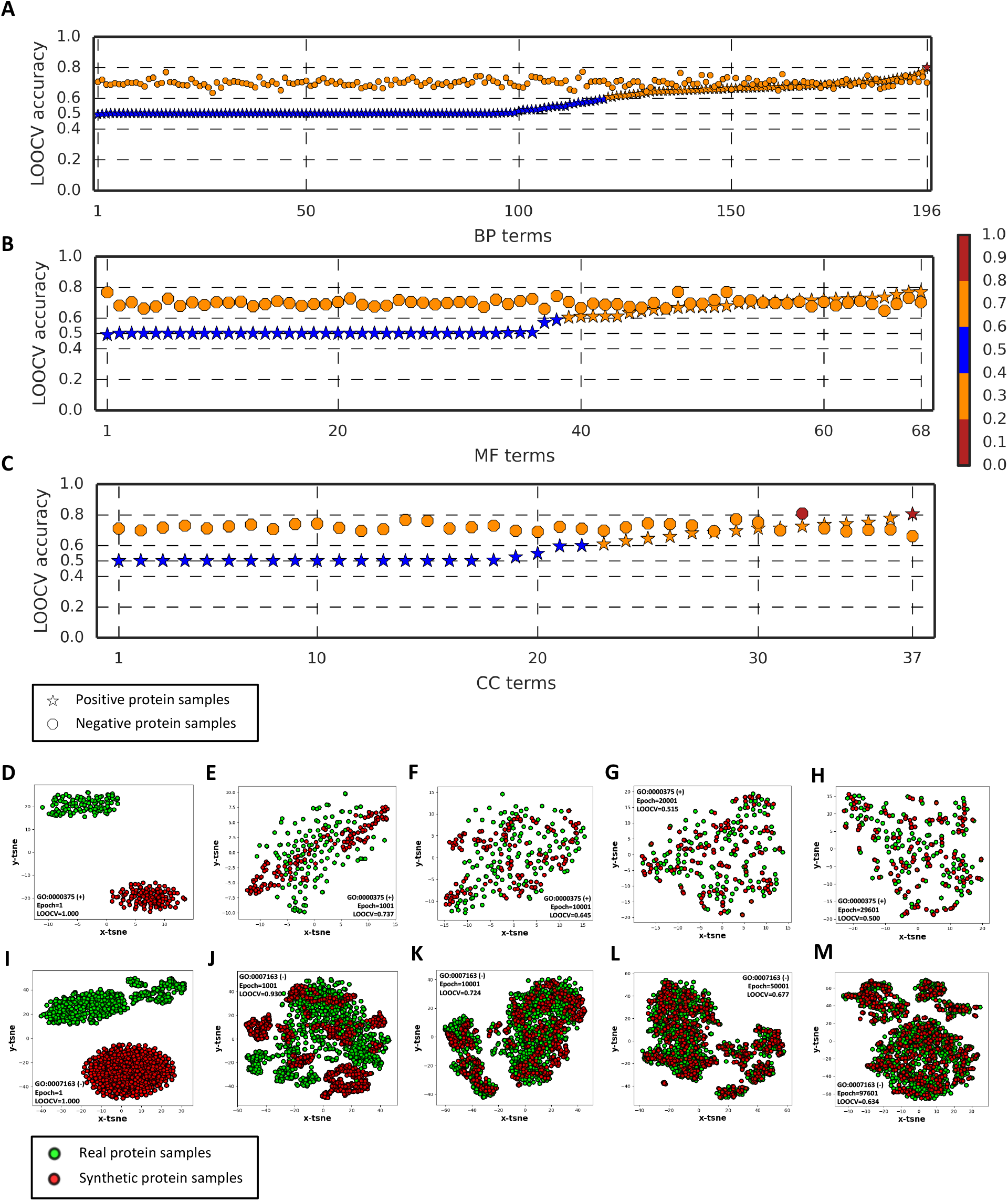
The Leave One Out Cross-Validation (LOOCV) accuracy of CTST obtained by the real and synthetic protein samples for GO terms from Biological Process (BP) (A), Molecular Function (MF) (B) and Cellular Component (CC) (C) domains; The *t*-SNE transformed 2D visualisation of real (green dots) and synthetic protein feature samples (red dots) obtained during different training epochs of FFPred-GAN by using positive and negative protein feature samples for BP terms GO:0000375 (D-H), GO:0007163 (I-M).

Due to the much higher diversity of negative feature samples, in that there are few ways of representing a positive but many ways of representing a negative case, accuracy for negative cases is lower, as might be expected. The LOOCV accuracies obtained by the synthetic negative protein feature samples range between 0.600 to 0.800 for all three domains of GO terms. The average LOOCV accuracies are 0.700, 0.698 and 0.720, respectively for BP, MF and CC domains. Analogously to the cases when training the synthetic positive protein feature samples, at the beginning of the FFPred-GAN training stage (i.e. after the 1^*st*^ epoch), both the real and synthetic negative samples for term GO:0007163 are obviously different, since both sets are distributed in different areas in Figure 2.I. After 1,001 epochs training, the distributions for both sets start to overlap, but the LOOCV accuracy of 0.930 is still far from optimal. After the 10,001^*st*^ epoch, the overlapping areas of both sets’ distributions become broader, with an improved LOOCV accuracy of 0.724. The training quality of FFPred-GAN continues to be improved even after 50,001 epochs of training, and finally the optimal negative synthetic protein feature samples are obtained after 97,601 epochs training, with an optimal LOOCV accuracy of 0.634. As shown in Figure 2.M, both green and red dots distribute in a similar area. This pattern is consistent when training FFPred-GAN for other two domains of GO terms, such as GO:0046872 and GO:0016020. As shown in Figure S1.O and S1.T, the real and synthetic negative protein feature samples for those two terms distribute in similar patterns after 52,201 and 49,001 epochs training, leading to the optimal LOOCV accuracies of 0.648 and 0.661, respectively.

### The synthetic protein feature samples generated by FFPred-GAN successfully improve the predictive accuracy of Drosophila function annotation using FFPred-fly

We evaluate the predictive power from using synthetic protein feature samples on the task of protein function prediction applied to *Drosophila*. We integrated the synthetic and real protein feature samples as the augmented training protein feature samples in 8 different ways, as shown on the first column of Table 1, where the rest of columns display the average MCC and AUROC values obtained by 3 different classification algorithms over individual domains of GO terms. In general, when using the Support Vector Machine (SVM) classification algorithm, 2 out of 8 different combinations of synthetic and real protein feature samples (i.e. Syn. Pos. + Real Pos. + Real Neg. and Syn. Pos. + Syn. Neg. + Real Pos. + Real Neg.) obtain higher accuracy on predicting all three domains of GO terms. The former obtains the overall highest accuracy on predicting the biological process (i.e. the average MCC of 0.345 and the average AUROC of 0.741) and molecular function (i.e. the average MCC of 0.664 and the average AUROC of 0.905) domains of protein function. It also obtains the same highest average AUROC value of 0.853 with the benchmark method for predicting the cellular component domain of protein function. Analogously, the latter obtains the overall highest average MCC value of 0.512 for predicting the cellular component domain of protein function, while it also obtains the same overall highest average AUROC value of 0.853 with the benchmark method.

**Table 1:** Summary of average MCC and AUROC values obtained by different combinations of training protein feature samples on 30% held-out testing samples by using different classification algorithms

| Instance Group | Support Vector Machine |  |  |  |  |  | *k*-Nearest Neighbours |  |  |  |  |  | Random Forests |  |  |  |  |  |
| --- | --- | --- | --- | --- | --- | --- | --- | --- | --- | --- | --- | --- | --- | --- | --- | --- | --- | --- |
|  | BP |  | MF |  | CC |  | BP |  | MF |  | CC |  | BP |  | MF |  | CC |  |
|  | MCC | AUROC | MCC | AUROC | MCC | AUROC | MCC | AUROC | MCC | AUROC | MCC | AUROC | MCC | AUROC | MCC | AUROC | MCC | AUROC |
| Syn. Pos. + Real Neg. | 0.182 | 0.687 | 0.468 | 0.868 | 0.317 | 0.806 | 0.276 | 0.614 | 0.555 | 0.746 | 0.397 | 0.671 | 0.001 | 0.541 | 0.000 | 0.586 | 0.005 | 0.590 |
| Syn. Neg. + Real Pos. | 0.179 | 0.700 | 0.394 | 0.871 | 0.338 | 0.820 | 0.273 | 0.637 | 0.544 | 0.779 | 0.443 | 0.720 | 0.022 | 0.608 | 0.086 | 0.744 | 0.050 | 0.715 |
| Syn. Pos. + Syn. Neg. | 0.232 | 0.691 | 0.485 | 0.857 | 0.402 | 0.808 | 0.251 | 0.603 | 0.507 | 0.725 | 0.397 | 0.667 | 0.130 | 0.633 | 0.360 | 0.779 | 0.284 | 0.719 |
| Syn. Pos. + Real Pos. + Real Neg. | **0.345\*** | **0.741\*** | **0.664\*** | **0.905\*** | **0.510** | **0.853\*** | **0.311** | 0.673 | **0.577** | 0.810 | 0.463 | 0.763 | 0.293 | 0.718 | 0.630 | 0.902 | 0.471 | 0.834 |
| Syn. Neg. + Real Pos. + Real Neg. | 0.302 | 0.726 | 0.653 | 0.901 | 0.465 | 0.850 | **0.307** | 0.643 | **0.588** | 0.788 | 0.464 | 0.724 | 0.316 | 0.731 | 0.627 | 0.901 | 0.475 | 0.833 |
| Syn. Pos. + Syn. Neg. + Real Pos. | 0.189 | 0.716 | 0.409 | 0.882 | 0.352 | 0.843 | 0.272 | 0.645 | 0.541 | 0.786 | 0.441 | 0.727 | 0.042 | 0.615 | 0.151 | 0.786 | 0.103 | 0.723 |
| Syn. Pos. + Syn. Neg. + Real Neg. | 0.187 | 0.690 | 0.491 | 0.871 | 0.328 | 0.807 | 0.264 | 0.598 | 0.529 | 0.727 | 0.387 | 0.653 | 0.008 | 0.560 | 0.018 | 0.658 | 0.044 | 0.631 |
| Syn. Pos. + Syn. Neg. + Real Pos. + Real Neg. | **0.344** | **0.735** | **0.662** | **0.903** | **0.512\*** | **0.853\*** | **0.310** | 0.651 | **0.590** | 0.798 | **0.466** | 0.726 | 0.323 | **0.740** | 0.619 | 0.902 | 0.471 | **0.852** |
| Real Pos. + Real Neg. (Benchmark) | 0.320 | 0.735 | 0.654 | 0.902 | 0.483 | <u>0.853*</u> | 0.301 | <u>0.698</u> | 0.575 | <u>0.850</u> | 0.465 | <u>0.775</u> | <u>0.333</u> | 0.734 | <u>0.631</u> | <u>0.905*</u> | <u>0.482</u> | 0.847 |

We further conduct pairwise comparisons between the best-performing combinations of protein feature samples and the benchmark protein feature samples, i.e. Syn. Pos. + Real Pos. + Real Neg. v.s. Real Pos. + Real Neg. for the biological process and molecular function domains, and Syn. Pos. + Syn. Neg. + Real Pos. + Real Neg. v.s. Real Pos. + Real Neg. for the cellular component domain of protein function. As shown in the scatter-plots in Figure 3, the majority of green dots drop above the diagonal, indicating higher MCC and AUROC values compared with the ones obtained by the benchmark training protein feature samples. In detail, using a combination of Syn. Pos. + Real Pos. + Real Neg., 106 out of 196 and 103 out of 196 biological process terms show improved results measured by MCC and AUROC values, respectively. This combination also gives higher MCC and AUROC values for 37 out of 68 (MCC) and 34 out of 68 (AUROC) molecular function terms. Analogously, the combination of Syn. Pos. + Syn. Neg. + Real Pos. + Real Neg. obtains higher MCC values for predicting 25 out of 37 cellular component terms. The pairwise Wilcoxon signed-rank tests further confirm that the augmented training protein feature samples obtain significantly higher MCC values for predicting all three domains of protein function (i.e. p-value of 5.86×10^-3^ for BP terms, p-value of 3.91 ×10^-2^ for MF terms, and p-value of 5.29×10^-3^ for CC terms). The AUROC values for predicting biological process domain of protein function are also significantly improved by the augmented training protein feature samples, due to the p-value of 3.12×10^-2^.

**Figure 3:**
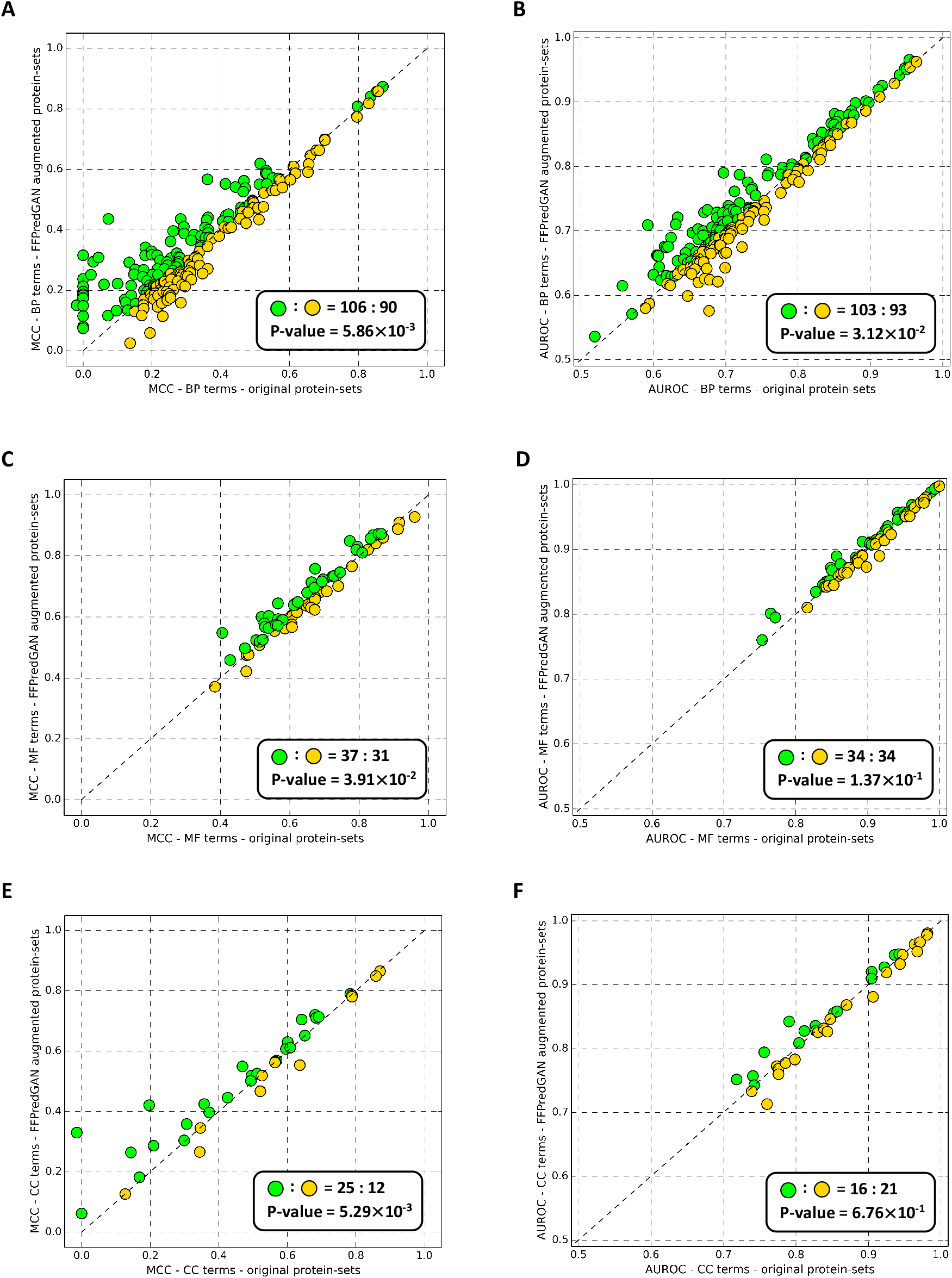
The scatter-plots about the MCC and AUROC values obtained by the optimal combination of real and synthetic protein samples for predicting three domains of GO terms by using the SVM classification algorithm.

The improved predictive accuracy is also observed on the augmented training protein feature samples using the *k*-nearest neighbours classification algorithm. Three types of combinations, i.e. Syn. Pos. + Real Pos. + Real Neg., Syn. Neg. + Real Pos. + Real Neg., and Syn. Pos. + Syn. Neg. + Real Pos. + Real Neg., obtain higher average MCC values than the benchmark training protein feature samples for predicting BP and MF terms, while the augmented protein feature samples consisting of Syn. Pos. + Syn. Neg. + Real Pos. + Real Neg. also obtain higher average MCC value for predicting CC terms. However, none of the augmented training protein feature samples obtains higher average AUROC value than the benchmark one. A similar pattern is also observed when using the random forests classification algorithm. The benchmark training protein feature samples obtain higher average MCC values than all other augmented training protein feature samples for predicting all three domains of protein function. But the augmented training protein feature samples consisting of Syn. Pos. + Syn. Neg. + Real Pos. + Real Neg. still obtain higher average AUROC values for predicting BP and CC terms..

## Discussion

### The optimal synthetic positive protein feature samples derive better SVM decision boundaries by augmenting the original training samples

The synthetic positive protein feature samples successfully improve the accuracy of predicting all three domains GO terms using a Support Vector Machine (SVM) classification algorithm. This fact suggests that the augmented training protein feature samples successfully derive better SVM decision boundaries. We further analyse the changes on the SVM decision boundaries with an example case of predicting the term GO:0034613 by using the original training protein feature samples and the synthetic positive protein feature samples augmented training samples respectively. The former leads to an MCC value of 0.073, whereas the latter leads to an MCC value of 0.436. We visualise the 2D distributions of both protein-sets by using their first two principle components, which are also used for training the SVM classifiers for visualising the corresponding 2D decision boundaries. As shown in Figures 4.A and 4.B, the blue dots denote the negative protein samples, while the red dots denote the positive protein samples. The white areas in the background denote the decision boundaries separating the blue and red areas where the negative and positive protein samples distributed. It is clear that the decision boundaries shown in both figures are different. The ones in Figure 4.A suggest that the SVM trained by the original protein samples successfully learned the boundaries that separate the protein samples with different labels in the centre of the figure. However, as shown in Figure 4.C, the boundaries learned by the original training protein-sets fail to separate the majority of negative and positive testing protein samples distributing on the right corner of the figure, where the majority of dots are in red. On the contrary, the SVM trained by the augmented training protein feature samples learned those decision boundaries that successfully separate the protein samples distributed on the right corner of the figure. As shown in Figure 4.D, when applying those decision boundaries on the testing protein feature samples, most of the red and blue dots on the right corner are successfully distinguished, leading to the increased MCC values.

**Figure 4:**
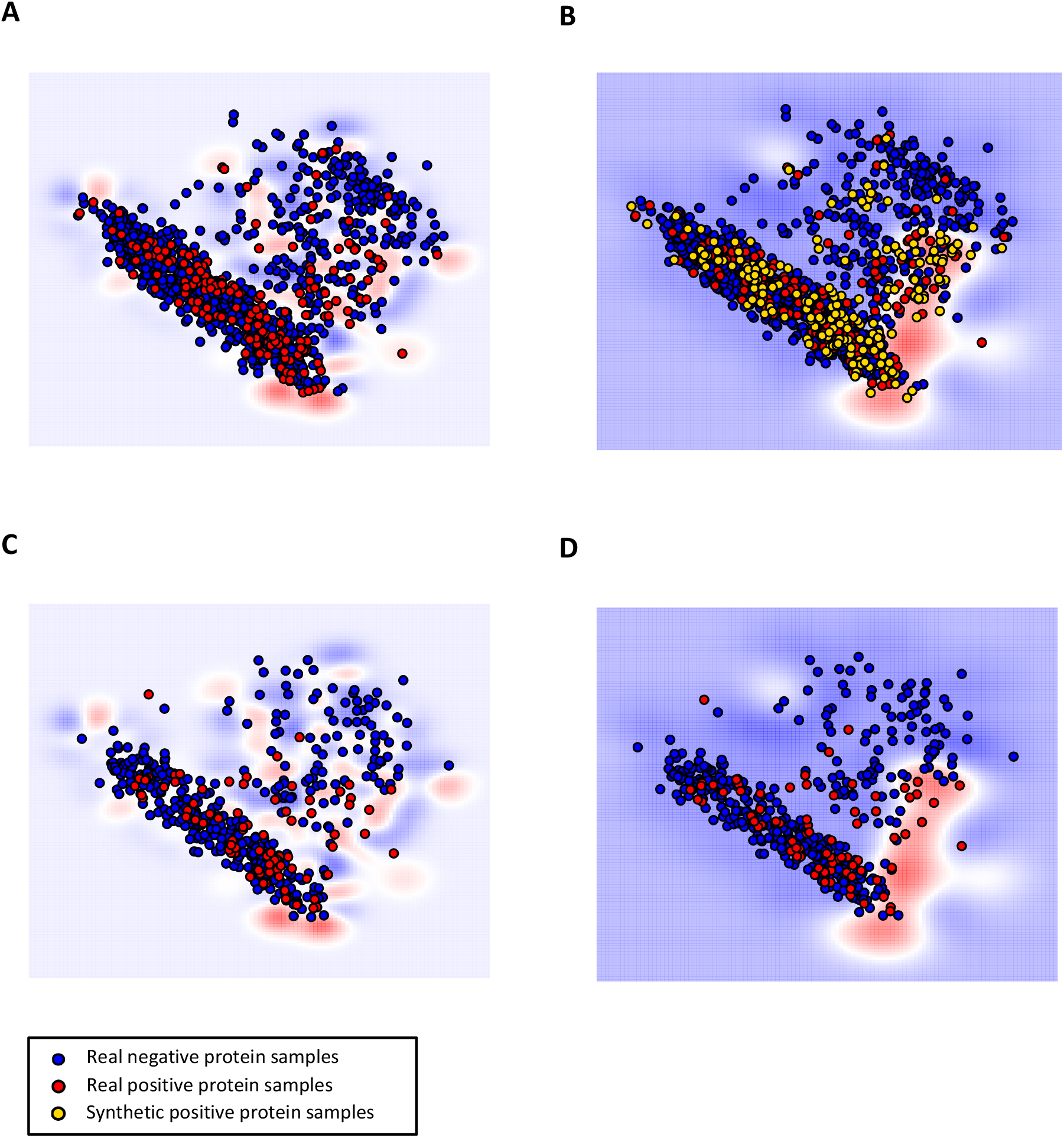
The 2D visualisations of the learned SVM decision boundaries by the original training protein feature samples (A) and the synthetic positive protein feature samples augmented training samples (B) for the term GO:0034613, and the distributions of corresponding testing protein feature samples with those two types of decision boundaries (C-D).

### FFPred-GAN can generate high-quality synthetic feature samples at reasonable computational cost

We further discuss the computational time cost (i.e. the actual running time obtained by using CPU-based PyTorch with a standard Linux computing cluster) and the training sample sizes (i.e. the number of training protein feature samples) for running FFPred-GAN to generate the optimal synthetic protein feature samples for individual GO terms. Figures 5.A and 5.B display the boxplots about the distributions of computational time and training samples size, respectively, while the complete information is reported in Table S1. In general, the computational time for generating optimal synthetic positive protein samples for the majority GO terms from all three individual domains (shown by the blue, golden and green boxes) is less than that of generating the optimal negative protein samples, as shown by the yellow, grey and orange boxes. The corresponding median values are 20,038.6 seconds (≈ 5.6 hours), 24,187.2 seconds (≈ 6.7 hours) and 20,973.8 seconds (≈ 5.8 hours) for generating the optimal positive synthetic protein samples respectively for BP, MF and CC domains of GO terms, while the median values for generating the optimal negative protein samples are 28.624.6 seconds (≈ 8.0 hours), 29,401.6 seconds (≈ 8.2 hours) and 113,777.8 seconds (≈ 31.6 hours) respectively for those three domains of GO terms. This fact is relevant with a pattern that the training samples sizes of positive proteins for the majority of GO terms are smaller than the negative ones, as shown in Figure 5.B, where the blue, golden and green boxes are located in lower positions than the yellow, grey and orange boxes. Analogously, the corresponding median values of sample sizes for those positive protein samples are 226.0, 234.0 and 238.0, respectively for BP, MF and CC domains of GO terms, while the median values of samples sizes for those negative protein samples are 873.0, 875.0 and 1680.0, respectively.

**Figure 5:**
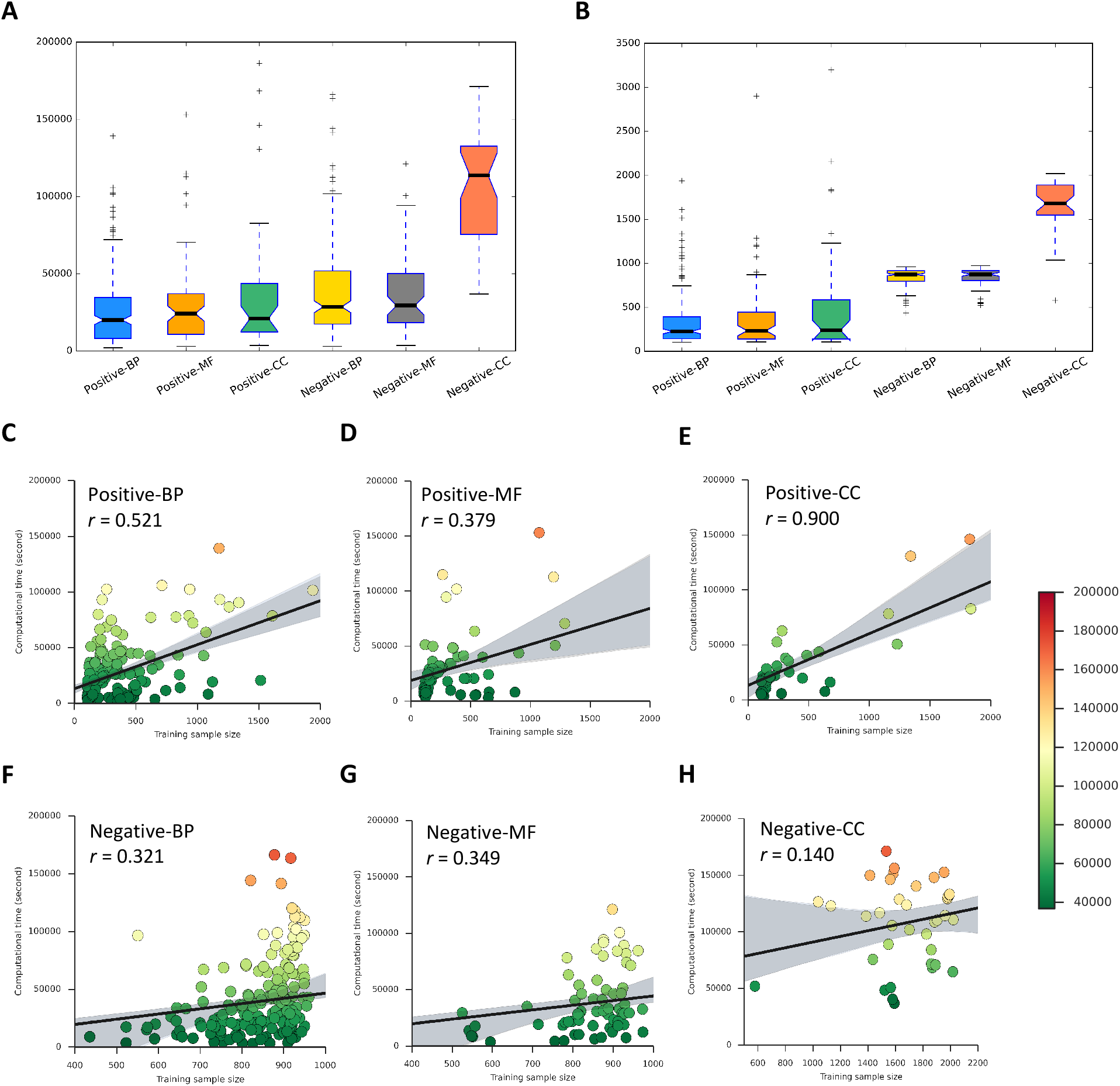
(A) The boxplot about the distributions of computational time on obtaining the optimal synthetic protein samples for different GO terms; (B) The boxplot about the distributions of sample sizes for different GO terms; (C-H) The scatter-plots of correlation coefficient values between the computational time and sample sizes for positive and negative protein samples of different domains of GO terms.

We then further calculate the Pearson correlation coefficient between the computational time and the training samples sizes, as shown by the scatter-plots in Figures 5.C – 5.H, where the *x*-axis denotes the values of sample size and the *y*-axis denotes the computational time. The correlation coefficient values *r* for positive protein samples are 0.521, 0.379 and 0.900 respectively for BP, MF and CC domains of GO terms, while the negative protein samples have the correlation coefficient values of 0.321, 0.349 and 0.140 respectively. Both the positive and negative protein samples from all three domains of GO terms all show positive correlation between the computational time and training samples size. This fact indicates that the larger samples size leads to longer training time of FFPred-GAN in order to obtain the optimal synthetic protein samples.

In this work, we have presented a novel generative adversarial networks-based method that successfully generates high-quality synthetic feature samples, which significantly improve the accuracy on predicting all three domains of GO terms through augmenting the original training data. Based on this same framework, there is significant scope to employ new GANs-based architectures, but more importantly, the same basic approach can be applied to other types of features used in function prediction, e.g. proteomics or gene expression data, which are often difficult or expensive to produce in large quantities. Finally, perhaps the most useful benefit of using GANs to augment data is that they can offer a powerful means to balance training sets in the usual situation of having many examples of proteins with one GO term label and very few of others. We hope to explore these applications in the future.

## Method details

### Generating synthetic protein feature samples with Wasserstein generative adversarial networks with gradient penalty

Wasserstein generative adversarial networks with gradient penalty [11] are a type of Generative Adversarial Networks (GANs) [8], which are well-known to be highly capable of learning highdimensional distributions from data samples. In general, conventional GANs are composed of two neural networks, i.e. the generator *G* and the discriminator (a.k.a. critic) *D*. The former takes the random Gaussian noise (a.k.a. the latent variables 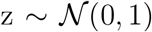) as the inputs to generate outputs that are considered as the synthetic samples. The latter takes the synthetic or real samples as the inputs to distinguish whether they are synthetic or not. In order to train the whole GANs, those two networks play a minimax two-player game, i.e. the generator aims to generate the synthetic samples as good as possible in order to fool the discriminator, whereas the discriminator aims to distinguish the real and synthetic samples as well as possible, as shown by Equation 1 (the minimax objective). Ideally, the GANs are successfully trained when those networks reach the Nash equilibrium, i.e. the generator is trained to optimally encode the actual distribution of target samples, while the discriminator is trained to optimally distinguish the real and synthetic samples. Usually, the weights of the generator are updated after several epochs of discriminator training. In essence, this process is equivalent to minimising the Jensen–Shannon (JS) or Kullback–Leibler (KL) divergences between the target distribution and the one encoded by the generator, given an optimal discriminator

Wasserstein GAN (WGAN) [10] is a well-known extension of conventional GANs. It adopts the earth-mover (Wasserstein) distance to replace the JS or KL divergences to avoid the vanishing gradient problem due to their natural limitation on handling non-overlapping distributions. In addition, WGAN adopts the weight clipping mechanism to enforce the 1-Lipchitz constraint for the critic w.r.t. the corresponding inputs. More recently, another extension of GANs was proposed, namely WGAN with Gradient Penalty (WGAN-GP) that further improves the training stability of WGAN by adopting the penalty mechanism on the norm of gradient of the critic. The objective of WGAN-GP is shown in Equation 2, where the left two terms denote the loss of critic, and the right term denotes the term of gradient penalty (i.e. ensure the *L*_2_ norm penalty being around 1.00).

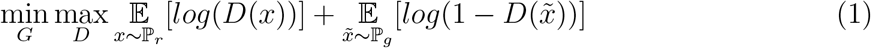

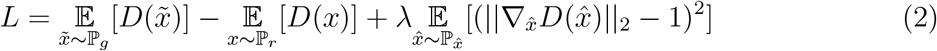

In this work, we use the generator of well-trained WGAN-GP models to generate synthetic samples. Each WGAN-GP model consists of a three hidden-layer generator and a three hidden-layer critic. The generator takes 258 dimensions of random Gaussian noise inputs and outputs 258 dimensions of synthetic samples. The ReLU activation function is adopted for all three hidden-layers (of 512 units each) followed by the output layer which adopts the tanh activation function. The critic network takes 258 dimensions of inputs (i.e. the real and synthetic protein feature samples), and uses the LeakyReLU activation function for all layers including three hidden-layers (of 86 units each). The Adam optimiser is used for training both generator and critic networks, with a learning rate of 1.00×10^-4^. The total number of epochs for training the WGAN-GP is 100,000, and the weights of the generator networks are updated after every 5 epochs of the critic training. The generated synthetic protein feature samples are saved after finishing every 200 epochs of WGAN-GP training for the purpose of down-stream quality assessment by using the classifier two-sample tests approach [33].

### Selecting optimal synthetic training protein feature samples by the classifier two-sample tests

FFPred-GAN evaluates and selects the optimal synthetic protein feature samples by using the Classifier Two-Sample Tests (CTST) [33] approach. The optimal synthetic protein feature samples are considered as those following the same distribution of the real (training) protein feature samples but being not identical to the real (training) ones. The CTST approach is an extension of conventional single variable-based statistical significance test methods (e.g. the Wilcoxon signed-rank test) to high-dimensional cases. More specifically, given two equal-sized sets of samples respectively following two distributions *P* and *Q*, the CTST consider to accept or reject a null hypothesis of *P* being not equal to *Q*. If the null hypothesis is rejected, the classification accuracy on predicting the binary labels of held-out samples will be near the chance-level (i.e. 50.0%). Therefore, in terms of a metric evaluating the quality of generated synthetic samples, a classification accuracy of 100.0% means the synthetic samples are of poor quality, due to the fact that the synthetic samples are significantly different to the real ones. Analogously, a classification accuracy of 0.0% also suggests poor quality of the synthetic samples, due to the fact that the real and synthetic samples appear identical, suggesting that the model is merely regenerating the training samples and has failed to generate diverse new synthetic samples.

In this work, we conduct the CTST by using the 1-nearest neighbour classification algorithm due to its simplicity on hyper-parameter tuning. The real and generated synthetic protein feature samples are merged as a union set of protein feature samples with being assigned the binary labels respectively, e.g. the label 1 for the real samples and the label 0 for the synthetic samples. The Leave One Out Cross-Validation (LOOCV) is used to obtain the classification accuracy of the CTST by using different synthetic protein feature samples during per 200 epochs of FFPred-GAN training. Finally, the synthetic protein feature samples that obtain the best LOOCV accuracy (i.e. closest to 50.0%) are selected as the optimal synthetic feature samples.

### Evaluating the predictive power of synthetic protein feature samples generated by FFPred-GAN for augmenting the original training samples

In this work, we use the same protein sets that were discussed in [5], i.e. 10,519 *Drosophila* proteins with 301 GO terms. The protein-set for each GO term was further split into the training and testing protein sets with a proportion of 7:3. The total 258 dimensions of protein sequence-derived biophysical features (in fact a mixture of distributions of different feature groups ranging from 11 to 50 dimensions) are used to describe the proteins, including information about the protein secondary structure, intrinsic disorder regions, signal peptides, etc. The full information about those feature groups is included in Table S2. The predictive power of the synthetic protein feature samples is evaluated by three different classification algorithms, i.e. support vector machine, *k*-nearest neighbours and random forests. A 5-fold cross validation-based grid-search is used for conducting the hyper-parameter optimisation for different classification algorithms. The detailed information about the hyper-parameter searching space is included in Table S3. The classification algorithms and grid-search procedure are implemented by using Scikit-learn [34]. The well-known Matthews Correlation Coefficient (MCC) and Area Under Receiver Operating Characteristic Curve (AUROC) are used to evaluate the predictive performance of FFPred-GAN. As shown in Equation 3, the MCC value is calculated by considering the true positive (TP), true negative (TN), false positive (FP) and false negative (FN) rates. Its ranges from −1 to 1, while a value of 0 means a random prediction and a value of 1 denotes perfect predictive accuracy. The AUROC value is another well-known metric on evaluating the accuracy of binary classification task. It is calculated by considering the true positive and false positive rates obtained by using different decision thresholds. The AUROC value ranges from 0 to 1, while a value of 0.5 indicates a random prediction and a value of 1.00 denotes perfect predictive accuracy.

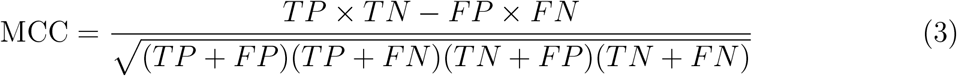

## Data and software availability

All data can be downloaded via http://bioinfadmin.cs.ucl.ac.uk/downloads/FFPredGAN The source-code can be downloaded via https://github.com/psipred/FFPredGAN

## Funding

C.W. and D.T.J. acknowledge funding from the European Research Council Advanced Grant ‘ProCovar’ (Project ID 695558) and the Biotechnology and Biological Sciences Research Council (BB/L002817/1). This work was supported by the Francis Crick Institute which receives its core funding from Cancer Research UK (FC001002), the UK Medical Research Council (FC001002), and the Wellcome Trust (FC001002).

## Acknowledgement

The authors acknowledge the members of the UCL Bioinformatics Group for valuable discussions. The authors also acknowledge the support of the high performance computing facility of the Department of Computer Science at the University College London.

## Declaration of interests

The authors declare no competing interests.

## Supporting File 1

### The Supporting Information includes

The *t*-SNE transformed 2D visualisation of real and optimal synthetic protein feature samples obtained during different training epochs of FFPred-GAN by using positive protein feature samples for molecular function term GO:0000981 and cellular component term GO: 0000785; and by using the negative protein feature samples for molecular function term GO: 0046872 and cellular component term GO:0016020. (Figure S1).

Summary of computational cost, training sample size and their correlation coefficient values. (Table S1).

Summary of protein amino acid sequence-derived biophysical feature groups information (Table S2).

Summary of hyper-parameters of different classification algorithms for conducting the 5-fold cross validation-based grid-search by using the Scikit-learn. (Table S3).

**Figure S1.**
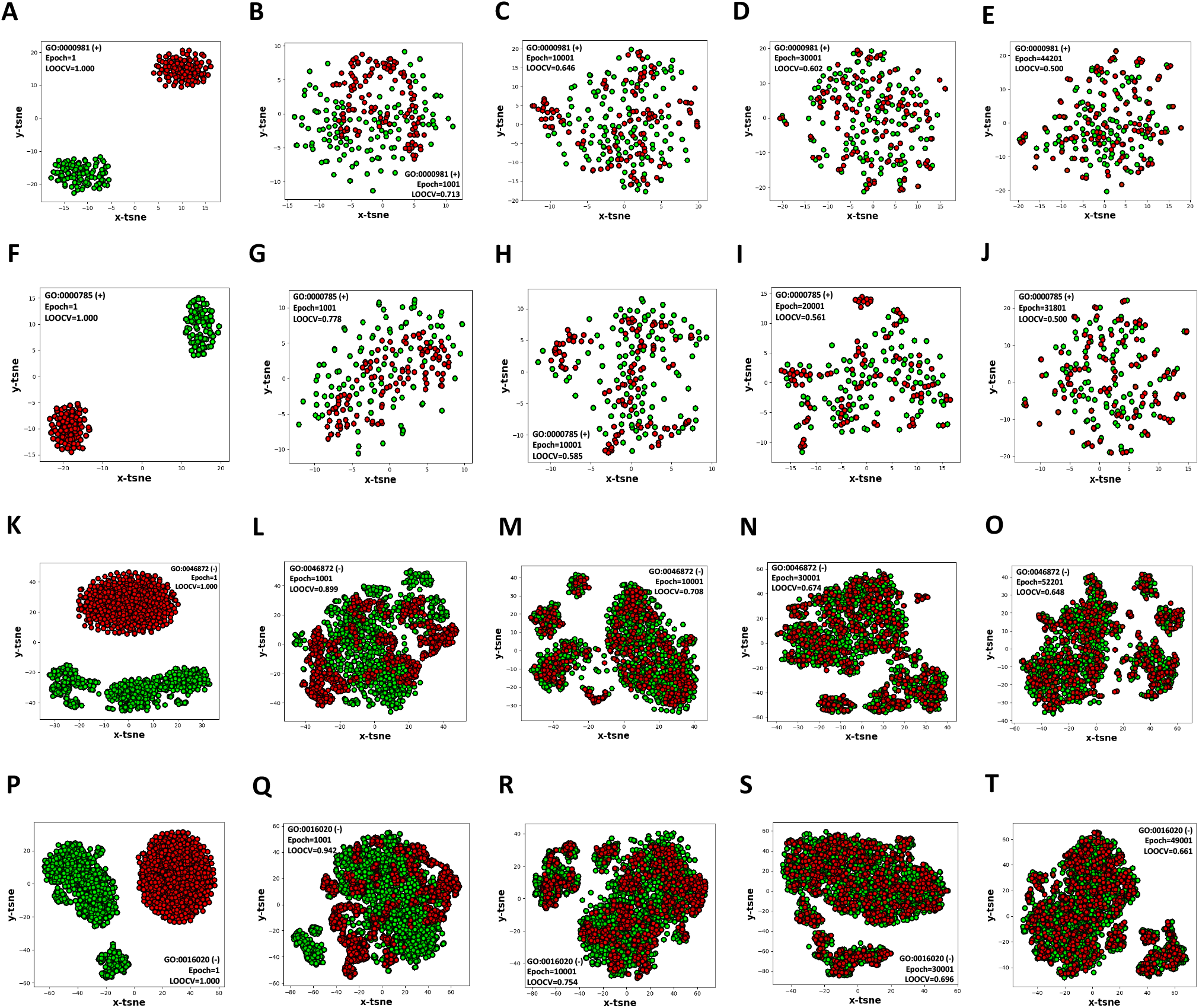
The *t*-SNE transformed 2D visualisation of real and optimal synthetic protein feature samples obtained during different training epochs of FFPred-GAN by using positive protein feature samples for molecular function term GO:0000981 (A-E) and cellular component term GO: 0000785 (F-J); and by using the negative protein feature samples for molecular function term GO: 0046872 (K-O) and cellular component term GO:0016020 (P-T).

**Table S1.** Summary of computational cost, training sample size and their correlation coefficient values

| Training Samples | Computational time cost |  |  |  |  |  |  |  |  |  | Training sample size |  |  |  |  | Correlation Coefficient |
| --- | --- | --- | --- | --- | --- | --- | --- | --- | --- | --- | --- | --- | --- | --- | --- | --- |
|  | Minimum |  | 1<sup>st</sup> Quartile |  | Median |  | 3<sup>rd</sup> Quartile |  | Maximum |  | Minimum | 1<sup>st</sup> Quartile | Median | 3<sup>rd</sup> Quartile | Maximum |  |
|  | Second | Hour | Second | Hour | Second | Hour | Second | Hour | Second | Hour |  |  |  |  |  |  |
| Positive-BP | 2073.0 | 0.6 | 8079.4 | 2.2 | 20038.6 | 5.6 | 34556.1 | 9.6 | 139210.2 | 38.7 | 105.0 | 143.8 | 226.0 | 392.5 | 1939.0 | 0.521 |
| Positive-MF | 3105.7 | 0.9 | 10887.2 | 3.0 | 24187.2 | 6.7 | 37066.0 | 10.3 | 153056.2 | 42.5 | 109.0 | 139.5 | 234.0 | 444.0 | 2901.0 | 0.379 |
| Positive-CC | 3520.0 | 1.0 | 12390.5 | 3.4 | 20973.8 | 5.8 | 43664.0 | 12.1 | 186317.1 | 51.8 | 106.0 | 140.0 | 238.0 | 584.0 | 3200.0 | 0.900 |
| Negative-BP | 2991.8 | 0.8 | 17456.5 | 4.8 | 28624.6 | 8.0 | 51873.3 | 14.4 | 166180.2 | 46.2 | 436.0 | 795.0 | 873.0 | 915.0 | 960.0 | 0.321 |
| Negative-MF | 3615.1 | 1.0 | 18286.3 | 5.1 | 29401.6 | 8.2 | 50182.8 | 13.9 | 121113.6 | 33.6 | 525.0 | 805.0 | 875.0 | 916.3 | 974.0 | 0.349 |
| Negative-CC | 36847.1 | 10.2 | 75593.5 | 21.0 | 113777.8 | 31.6 | 132606.9 | 36.8 | 171204.0 | 47.6 | 579.0 | 1548.0 | 1680.0 | 1891.0 | 2020.0 | 0.140 |

**Table S2.** Summary of protein sequence-derived biophysical feature groups information.

| Feature names | Dimensions |
| --- | --- |
| Secondary structure | 49 |
| Transmembrane segments | 15 |
| Amino acid composition | 20 |
| Intrinsically disordered regions | 18 |
| Signal peptides | 8 |
| Subcellular localization | 12 |
| Sequence features | 17 |
| PEST regions | 12 |
| Low complexity regions | 12 |
| Coiled coils | 12 |
| N-linked glycosylation sites | 11 |
| O-GalNAc-glycosylation sites | 22 |
| Phosphorylation sites | 50 |

**Table S3.**
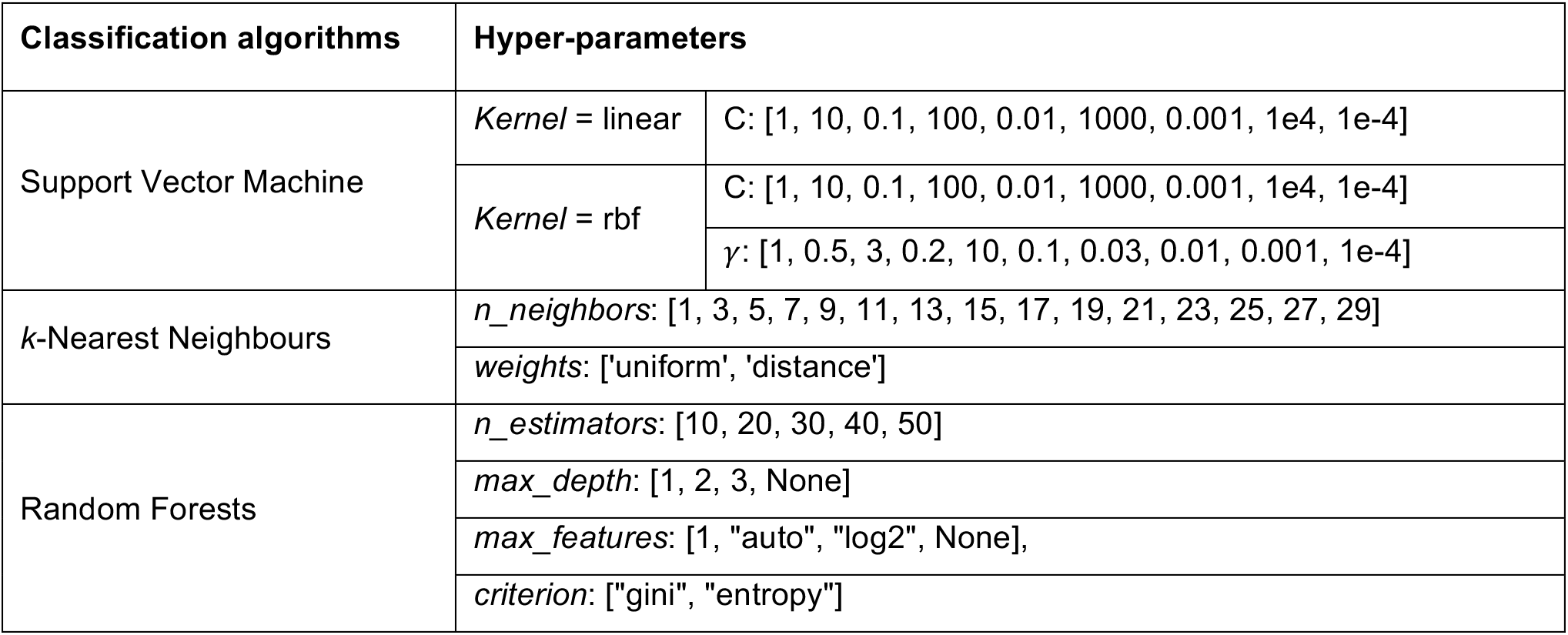
Summary of hyper-parameters of different classification algorithms for conducting the 5-fold cross validation-based grid-search by using the Scikit-learn.

